# The gut microbiome of the healthy population in Kolkata, India, is a reservoir of antimicrobial resistance genes emphasizing the need of enforcing antimicrobial stewardship

**DOI:** 10.1101/2023.07.17.549428

**Authors:** Rituparna De, Suman Kanungo, Asish Kumar Mukhopadhyay, Shanta Dutta

**Affiliations:** Division of Bacteriology, National Institute of Cholera and Enteric Diseases, Kolkata, India; Division of Epidemiology, National Institute of Cholera and Enteric Diseases, Kolkata, India

**Keywords:** Antimicrobial resistance, gut microbiome, community, antimicrobial stewardship, SDG, diarrhea

## Abstract

Antimicrobial resistance (AMR) alleviation warrants antimicrobial stewardship (AS) entailing indispensability of epidemiological surveillance. We undertook a small-scale surveillance in Kolkata to detect the presence of antimicrobial resistance genes (ARGs) in the healthy gut microbiome. We found that it was a reservoir of ARGs against common antibiotics. A targeted PCR and sequencing-based ARGs detection against tetracyclines, macrolides, trimethoprim, sulfamethoxazole, aminoglycosides, amphenicol and mobile genetic element (MGE) markers was deployed in twenty-five fecal samples. Relative abundance and frequency of ARGs was calculated. We detected markers against all these classes of antibiotics. 100% samples carried aminoglycoside resistance marker and *int1U*. A comparison with our previously published diarrheal resistome from the same spatial and temporal frame revealed that a higher diversity of ARGs were detected in the community and a higher rate of isolation of *tetC, msrA, tmp* and *sul-2* was found. The presence of common markers in the two cohorts proves that the gut microbiome has been contaminated with ARGs and which are being disseminated among different ecosystems. This is an issue of discerning concern for public health. The study raises an alarming picture of the AMR crisis in low-middle and emergent economies. It emphasizes the strict enforcement of AS in the community.

## Introduction

Antimicrobial resistance (AMR) is a serious public health concern challenging the success of antibiotics in treating infections and consequently leading to higher mortality and morbidity (1,2). AMR in bacteria arises due to the presence of antimicrobial resistance genes (ARGs) in the genome of bacteria (3). These ARGs encode proteins which impair the activity of antibiotics and are disseminated by HGT (Horizontal Gene Transfer) with the help of MGEs (mobile genetic elements) (3). Acquisition of ARGs is stimulated under circumstances which induce stress in the environment like the presence of antibiotics and other agents (3) and also produce persistor phenotypes (1). These ensure a cycle of ARG dissemination which is difficult to intercept (3). The microbiome has been found to be a conducive milieu facilitating ARG dissemination (3). Epidemiological surveillance as a routine measure of detecting the presence and diversity of ARGs is a crucial module of antimicrobial stewardship (AS) which serves as an early warning system to prevent devastating outcomes in public health (4). In this context, we conducted a pilot study to detect the presence of ARGs against common antibiotics in the gut microbiome of the community. Accordingly, PCR amplification was carried out using specific primers for detecting common ARGs, serving as markers of resistance against the major classes of antibiotics, namely, aminoglycosides, tetracyclines, macrolides, trimethoprim, sulfamethoxazole, amphenicol and an MGE, integron Class 1, 2 and 4. We tested the relative abundance of these ARGs and also the frequency of occurrence of resistance marker against these antibiotic classes. The study revealed that these ARGs are present in variable proportions in the healthy community gut microbiome. Therefore, the healthy gut microbiome is a potential reservoir of ARGs and a source of dissemination. This kind of epidemiological surveillance would help to screen the population for the presence and diversity of ARGs and help in the management of AMR.

## Materials and Methods

### Collection of specimen

25 fecal samples (CS1-CS25) were collected from ward numbers 58 and 59, both under Kolkata Municipal Corporation after obtaining informed consent from the donors on approval by the Scientific Advisory Committee (SAC) and the Institutional Ethics Committee (IEC). Donors were selected based on exclusion criteria of not reporting any infections nor chronic conditions, not on routine medication, not undergone antibiotic therapy in the preceeding 6 months before the date of collection of specimen.

### Molecular detection of ARGs

DNA was extracted following the GITC method (5). PCR amplification of the 16S rRNA conserved regions C1-C9 followed by PCR for the detection of the ARGs, *tetA, tetB, tetC, tetD, tetE, tetM, ermB, mefA, msrA, mphA, dfr1, tmp, dhrfr1, dfrA12, dfrA15, dfrA27, sul2, strA, strB, aadA1, aac(3), floR, int1U, int2U, int4U* encoding resistance to aminoglycosides, tetracyclines, macrolides, trimethoprim, sulfamethoxazole, amphenicol and an MGE, integron Class 1, 2 and 4. The presence of the ARG detected by positive PCR reaction was further confirmed by Sanger Sequencing. The method for molecular typing and sequences of primers used in the study were the same as described in our previous study (6).

Relative abundance of the different ARGs and frequency of occurrence of resistance against each class of antibiotic were calculated following the formulae used in our previous study (6). Relative abundance is the number of samples in which each ARG was detected expressed as percentage while the frequency against each class of antibiotic was expressed as the percentage of the number of samples in which at least one marker for each class was detected.

## Results

Table 1 represents the individual markers encoding resistance to aminoglycosides, tetracyclines, macrolides, trimethoprim, sulfamethoxazole and integron Class 1 marker which were detected and occurred in low, moderate and high abundance of up to 92% for *strA*, 96% for *sul-2* and *tetBM* respectively and 100% for *int1U*. Table 2 shows the ARG profile of CS1-25 for those ARGs which were detected in at least one sample. The amphenicol resistance marker was not detected. The rate of isolation of aminoglycoside resistance and integron class 1 marker was 100% each. Figures 1 and 2 represent relative abundance and frequency respectively.

**Table 1.** Relative abundance of ARGs in 25 fecal samples.

| No. of samples | ARG | % samples | Class Of antibiotic |
| --- | --- | --- | --- |
| 23 | *strA* | 92 | Aminoglycoside |
| 19 | *strB* | 76 | Aminoglycoside |
| 7 | *ant* | 28 | Aminoglycoside |
| 9 | *mphA* | 36 | Macrolide |
| 14 | *mefA* | 56 | Macrolide |
| 4 | *msrA* | 16 | Macrolide |
| 18 | *tetA* | 72 | Tetracycline |
| 24 | *tetB* | 96 | Tetracycline |
| 17 | *tetC* | 68 | Tetracycline |
| 24 | *tetM* | 96 | Tetracycline |
| 24 | *sul-2* | 96 | Sulfamethoxazole |
| 1 | *tmp(dfr18)* | 4 | trimethoprim |
| 16 | *aadA1* | 64 | trimethoprim |
| 11 | *dfr1 (dfr18)* | 44 | trimethoprim |
| 25 | *int1U* | 100 | Integron class 1 |

**Table 2.**
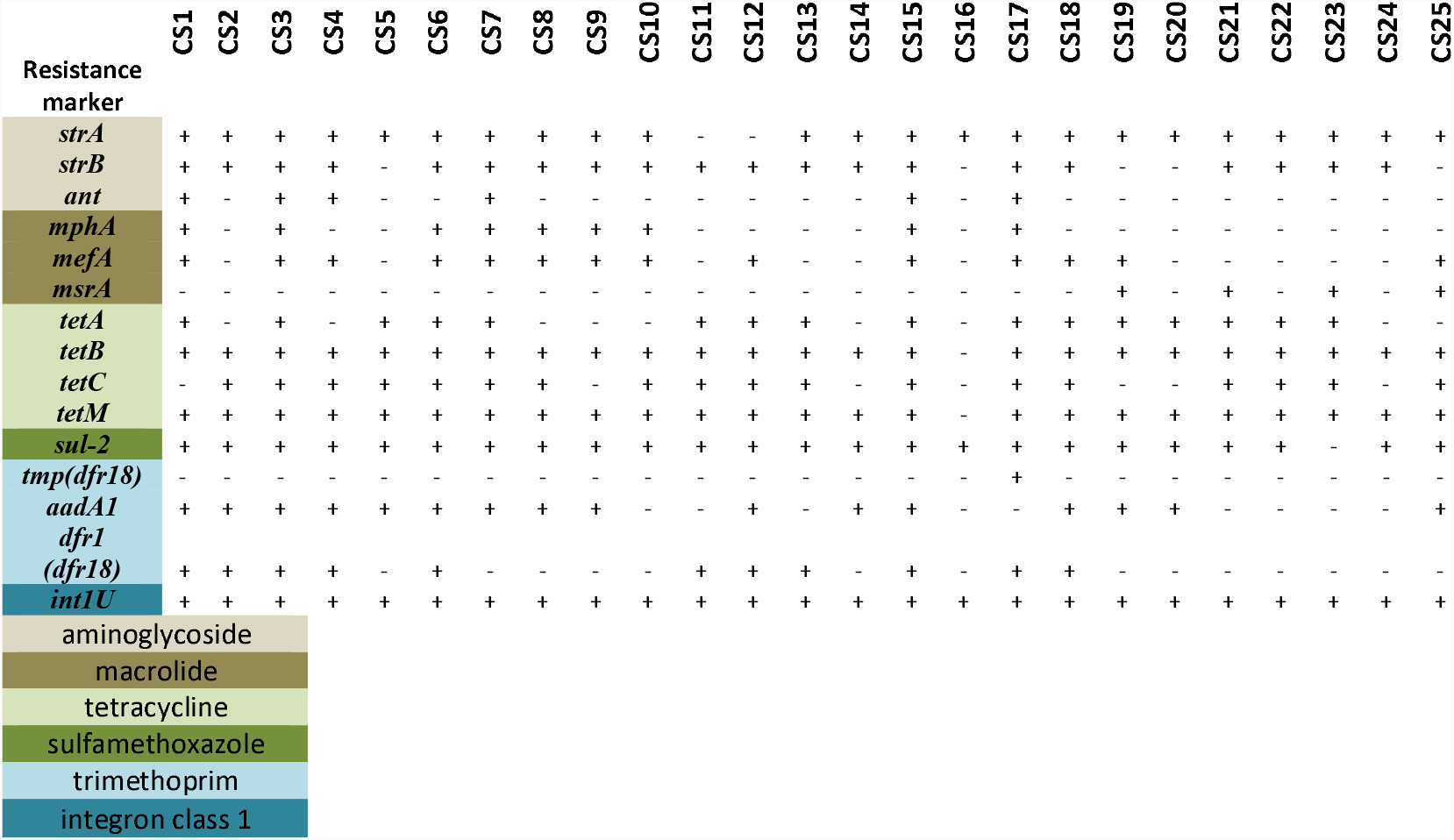
Table showing the results of PCR amplification of ARGs detected in CS1-CS25.

**Figure 1.**
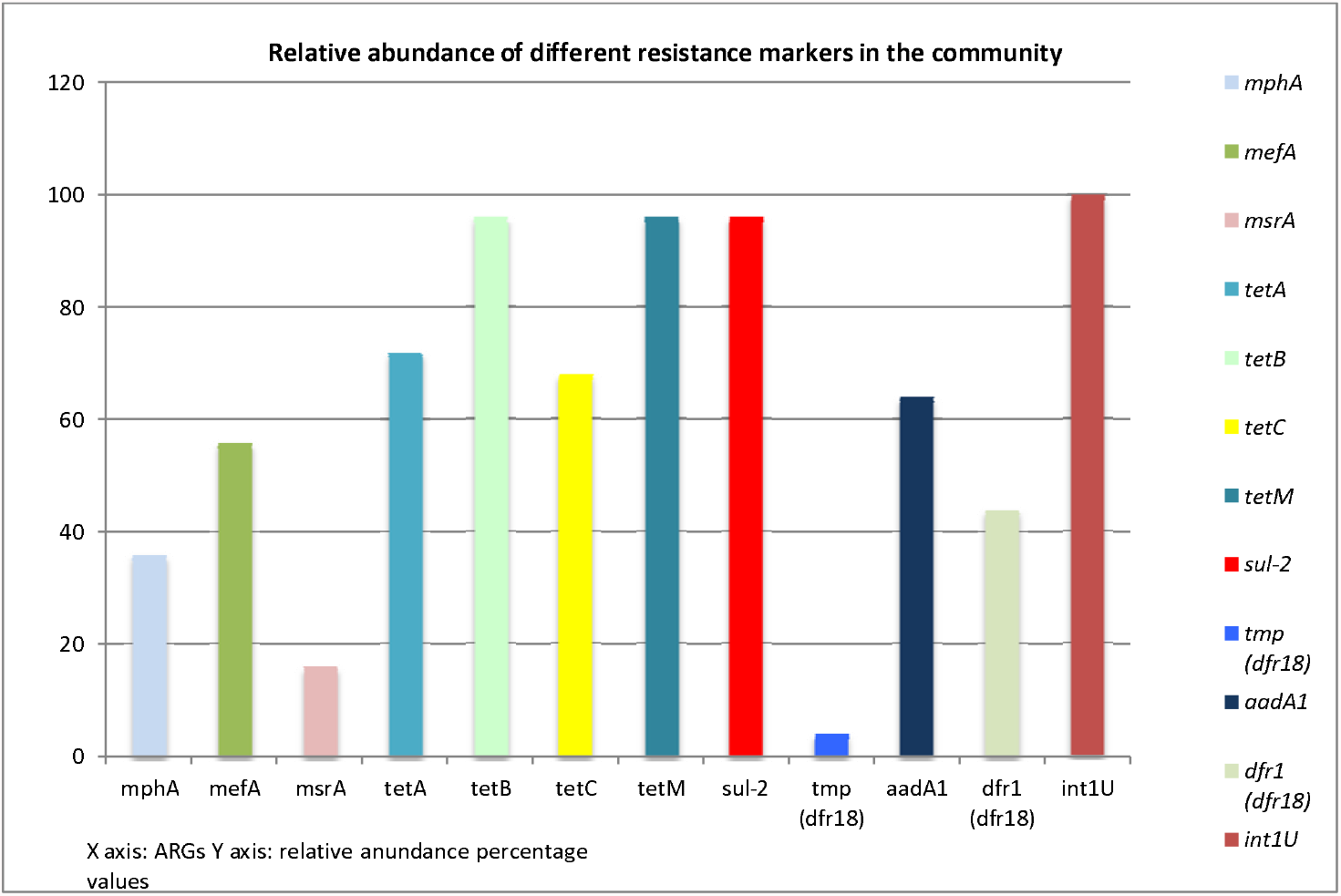
Bar-diagram showing the relative abundance of genetic markers of resistance against common antibiotics

**Figure 2.**
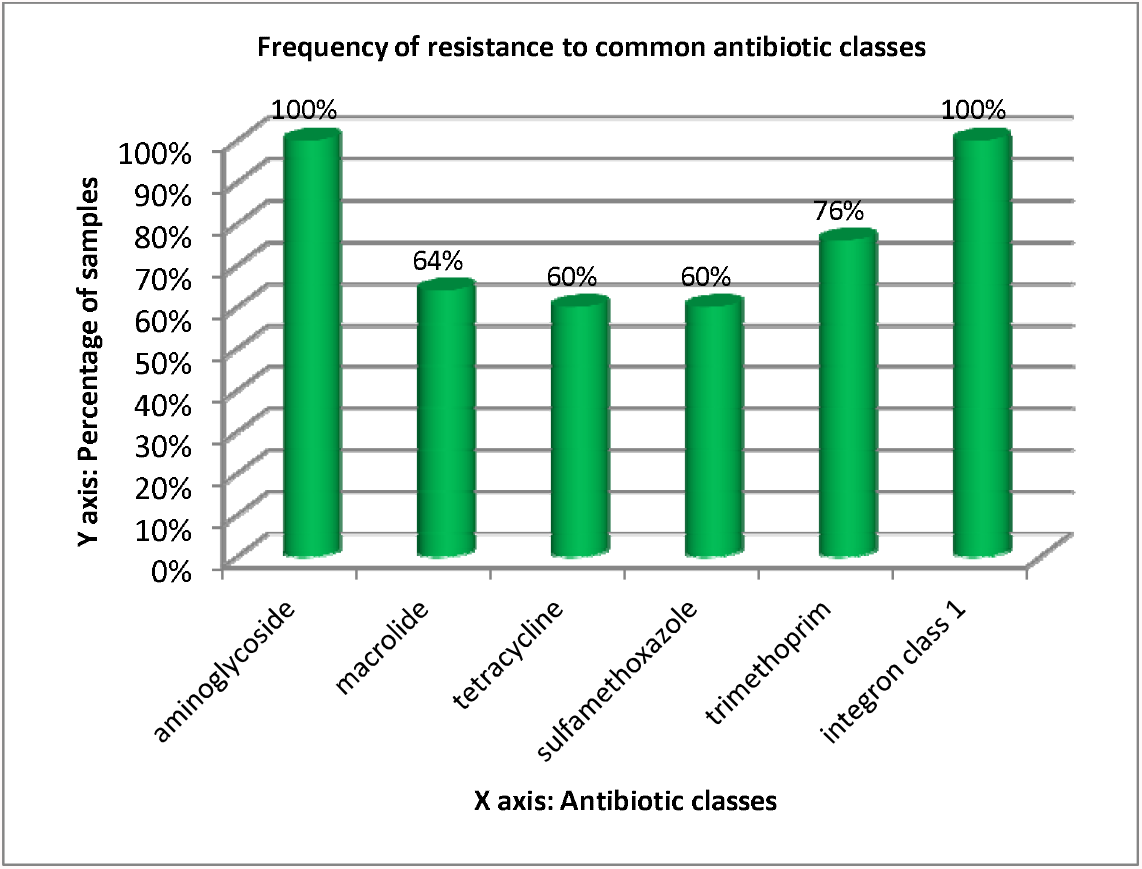
Bar-diagram showing the frequency of resistance to common antibiotics in the 25 community samples CS1-CS25

## Discussion

AMR is a major contributor of enhanced infectious disease mortality and morbidity due to MDR (multidrug resistant) and XDR (extensively drug resistant) pathogens recalcitrant to effective treatment (2).

It has been recognized as an urgent public health and economic challenge and is one of the foci of Sustainable Development Goals (SDGs) of WHO (7-8). The objective of diminishing the hazard of AMR had called for antimicrobial stewardship as part of the One Health goals of WHO (9-10).

The present study has been conducted in this context to perceive the threat of AMR in economically strapped settings, like the volunteers selected from ward number 58 and 59 of Kolkata in India represented by a population which is under the continuous threat of infectious diseases for which antibiotic therapy is indispensible (8, 11-14). At the same time, the overuse and misuse of antibiotics in this setting has been identified as a major contributory factor to the aggravating burden of AMR (11, 15). Consequently, microbial communities in the gut microbiome of the urban community have acquired ARGs which encode resistance to empirical antibiotics administered for common infections like diarrheal diseases (6). This has been revealed in the present study from the isolation of high abundance of individual ARGs and the frequency of isolation of markers of resistance. We compared ARG profile of community samples with those of the diarrheal patients reported in our previous published study (6). Comparatively, we found a higher abundance of *tetC* and *sul-2* genes of 68% and 96% in the community against 4% and 76% in diarrheal patients respectively from comparable demography and screened during the same time span 2019-2020. *msrA* and *tmp* genes were exclusively detected in the community. The findings from this study can be used to predict the pattern of AMR in common pathogens and may also guide the development of screening guidelines for antibiotic susceptibility testing in common pathogens. It also warrants strict implementation of antimicrobial stewardship to alleviate the burden of AMR to prevent arriving at the precipice when MDR infections will be completely untreatable.

## Acknowledgement

RD is grateful to Dr. G. Balakrish Nair and Dr. Swarup Sarkar for their motivation and help with the logistics for the project.

## References

1. Huemer M, Mairpady Shambat S, Brugger SD, Zinkernagel AS. Antibiotic resistance and persistence-Implications for human health and treatment perspectives. EMBO Rep. 2020 Dec 3;21(12):e51034. doi: 10.15252/embr.202051034.

2. Antimicrobial Resistance Collaborators. Global burden of bacterial antimicrobial resistance in 2019: a systematic analysis. Lancet. 2022 Feb 12;399(10325):629–655. doi: 10.1016/S0140-6736(21)02724-0.

3. De R (2021) Mobile Genetic Elements of Vibrio cholerae and the Evolution of Its Antimicrobial Resistance. Front. Trop. Dis 2:691604. doi: 10.3389/fitd.2021.691604

4. Cox JA, Vlieghe E, Mendelson M, Wertheim H, Ndegwa L, Villegas MV, Gould I, Levy Hara G. Antibiotic stewardship in low- and middle-income countries: the same but different? Clin Microbiol Infect. 2017 Nov;23(11):812–818. doi: 10.1016/j.cmi.2017.07.010.

5. De, Rituparna, Asish Kumar Mukhopadhyay, and Shanta Dutta. 2020. “Metagenomic Analysis of Gut Microbiome and Resistome of Diarrheal Fecal Samples from Kolkata, India, Reveals the Core and Variable Microbiota Including Signatures of Microbial Dark Matter.” Gut Pathogens 12 (1). https://doi.org/10.1186/s13099-020-00371-8.

6. De R, Mukhopadhyay AK, Dutta S. Molecular Analysis of Selected Resistance Determinants in Diarrheal Fecal Samples Collected From Kolkata, India Reveals an Abundance of Resistance Genes and the Potential Role of the Microbiota in Its Dissemination. Front Public Health. 2020 Mar 11;8:61. doi: 10.3389/fpubh.2020.00061.

7. https://www.who.int/news/item/22-06-2023-who-outlines-40-research-priorities-on-antimicrobial-resistance (Last accessed July 6, 2023)

8. https://www.who.int/news/item/24-03-2023-who-releases-priorities-for-research-and-development-of-age-appropriate-antibiotics (Last accessed July 6, 2023)

9. https://www.who.int/publications/i/item/9789241515481 (Last accessed July 6, 2023)

10. https://www.who.int/news/item/30-05-2018-international-partnership-to-address-human-animal-environment-health-risks-gets-a-boost (Last accessed July 6, 2023)

11. Laxminarayan R, Chaudhury RR. Antibiotic Resistance in India: Drivers and Opportunities for Action. PLoS Med. 2016 Mar 2;13(3):e1001974. doi: 10.1371/journal.pmed.1001974.

12. Menon GR, Singh L, Sharma P, Yadav P, Sharma S, Kalaskar S, Singh H, Adinarayanan S, Joshua V, Kulothungan V, Yadav J, Watson LK, Fadel SA, Suraweera W, Rao MVV, Dhaliwal RS, Begum R, Sati P, Jamison DT, Jha P. National Burden Estimates of healthy life lost in India, 2017: an analysis using direct mortality data and indirect disability data. Lancet Glob Health. 2019 Dec;7(12):e1675–e1684. doi: 10.1016/S2214-109X(19)30451-6.

13. Fuhrmeister ER, Harvey AP, Nadimpalli ML, Gallandat K, Ambelu A, Arnold BF, Brown J, Cumming O, Earl AM, Kang G, Kariuki S, Levy K, Pinto Jimenez CE, Swarthout JM, Trueba G, Tsukayama P, Worby CJ, Pickering AJ. Evaluating the relationship between community water and sanitation access and the global burden of antibiotic resistance: an ecological study. Lancet Microbe. 2023 Jun 30:S2666-5247(23)00137-4. doi: 10.1016/S2666-5247(23)00137-4.

14. Walia, K., & Gangakhedkar, R. R. (2021). Infectious disease blocks in district hospitals to augment India’s resolve to contain antimicrobial resistance. The Indian journal of medical research, 153(4), 416–420. https://doi.org/10.4103/ijmr.IJMR_4031_20

15. Tamhankar AJ, Karnik SS, Stålsby Lundborg C. Determinants of Antibiotic Consumption - Development of a Model using Partial Least Squares Regression based on Data from India. Sci Rep. 2018 Apr 23;8(1):6421. doi: 10.1038/s41598-018-24883-1.

